# BactMAP: an R package for integrating, analyzing and visualizing bacterial microscopy data

**DOI:** 10.1101/728782

**Authors:** Renske van Raaphorst, Morten Kjos, Jan-Willem Veening

**Affiliations:** Department of Fundamental Microbiology, Faculty of Biology and Medicine, University of Lausanne, CH-1015 Lausanne, Switzerland; Molecular Genetics Group, Groningen Biomolecular Sciences and Biotechnology Institute, Centre for Synthetic Biology, University of Groningen, 9747 AG Groningen, The Netherlands; Faculty of Chemistry, Biotechnology and Food Science, Norwegian University of Life Sciences, N-1432 Ås, Norway

**Keywords:** Image analysis, Bacterial cell biology, Single cell analysis, *Streptococcus pneumoniae*, *Staphylococcus aureus*, *Bacillus subtilis*, DNA replication, Chromosome segregation, Rtools

## Abstract

High-throughput analyses of single-cell microscopy data is a critical tool within the field of bacterial cell biology. Several programs have been developed to specifically segment bacterial cells from phase-contrast images. Together with spot and object detection algorithms, these programs offer powerful approaches to quantify observations from microscopy data, ranging from cell-to-cell genealogy to localization and movement of proteins. Most segmentation programs contain specific post-processing and plotting options, but these options vary between programs and possibilities to optimize or alter the outputs are often limited. Therefore, we developed BactMAP (<u>Bac</u>terial <u>t</u>oolbox for <u>M</u>icroscopy <u>A</u>nalysis & <u>P</u>lotting), a software package that allows researchers to transform cell segmentation and spot detection data generated by different programs automatically into various plots. Furthermore, BactMAP makes it possible to perform custom analyses and change the layout of the output. Because BactMAP works independently of segmentation and detection programs, inputs from different sources can be compared within the same analysis pipeline. BactMAP runs in R, which enables the use of advanced statistical analysis tools as well as easily adjustable plot graphics in every operating system. Using BactMAP we visualize key cell cycle parameters in *Bacillus subtilis* and *Staphylococcus aureus*, and demonstrate that the DNA replication forks in *Streptococcus pneumoniae* dissociate and associate before splitting of the cell, after the Z-ring is formed at the new quarter positions. BactMAP is available from https://veeninglab.com/bactmap.

## Introduction

Segmentation tools for analyses of phase contrast microscopy images of bacterial cells are improving rapidly. Over a decade ago the first software packages became available, making it possible to track cells (semi-) automatically using programs like Bacterial Home Vision (BHV), CellProfiler, Schnitzcells and MicrobeTracker (Stewart *et al*., 2005; Lamprecht *et al*., 2007; Sliusarenko *et al*., 2011; Young *et al*., 2011). However, using these tools and transforming the data could be time-consuming and still heavily relied on manual adjustments. Recent years, more user-friendly and quicker tools have been developed to segment phase-contrast images or fluorescence images, making cell segmentation a standard tool for bacterial microscopy analysis. With the creation of datasets of cell shape, size, growth and internal fluorescent signal information so easily at hand, the next challenge is to gain biologically useful insights.

Most of the current popular cell segmentation programs include basic tools for exploring the data in the form of histograms or fluorescence profiles. Oufti (Paintdakhi *et al*., 2016), ObjectJ (Vischer *et al*., 2015) and MicrobeJ (Ducret *et al*., 2016) have excellent tools for exploring several characteristics of the dataset. Oufti has a few fast time-lapse plotting and exploring tools, while MicrobeJ gives the user the possibility to plot characteristics of sub-groups of cells in many different ways. Within ObjectJ it is relatively easy to add custom measurements to the analysis and make quick visualizations. Supersegger (Stylianidou *et al*., 2016) includes a set of plotting and filtering tools inspired by flow cytometry analysis (Cass *et al*., 2017).

While the options for analysis within these programs are improving and expanding, there are limitations to having the analysis toolbox and the detection tool within the same program. After exploring the quality of the detection and the general distribution of the data by for instance fluorescence histograms and dot plots of the cell size, more in-depth custom analysis will often be necessary. Apart from that, multiple datasets containing data from different conditions or replicates will often need to be combined before analyzing them altogether. This is not always straightforward as the different software packages are mostly built to directly proceed from segmentation and detection. Finally, with so many new and excellent tools at hand for single molecule tracking, cell segmentation and cell shape identification, it can be very useful to combine datasets generated from different programs or to compare the output of similar programs to see which one is optimal for different purposes.

Therefore we created BactMAP (<u>B</u>acterial <u>t</u>oolbox for <u>M</u>icroscopy <u>A</u>nalysis & <u>P</u>lotting), an R package (https://www.r-project.org/), which acts as a funnel for segmentation and fluorescence detection data (Fig. 1). BactMAP orders the input data in a standard way, filtering out the most-used information but keeping all original data attached. It is possible to gather data from different sources and combine them, visualize the data and perform custom analysis using the BactMAP toolbox together with the vast amount of tools already available in R. BactMAP, its code and a user guide is available at https://veeninglab.com/bactmap.

**Figure 1.**
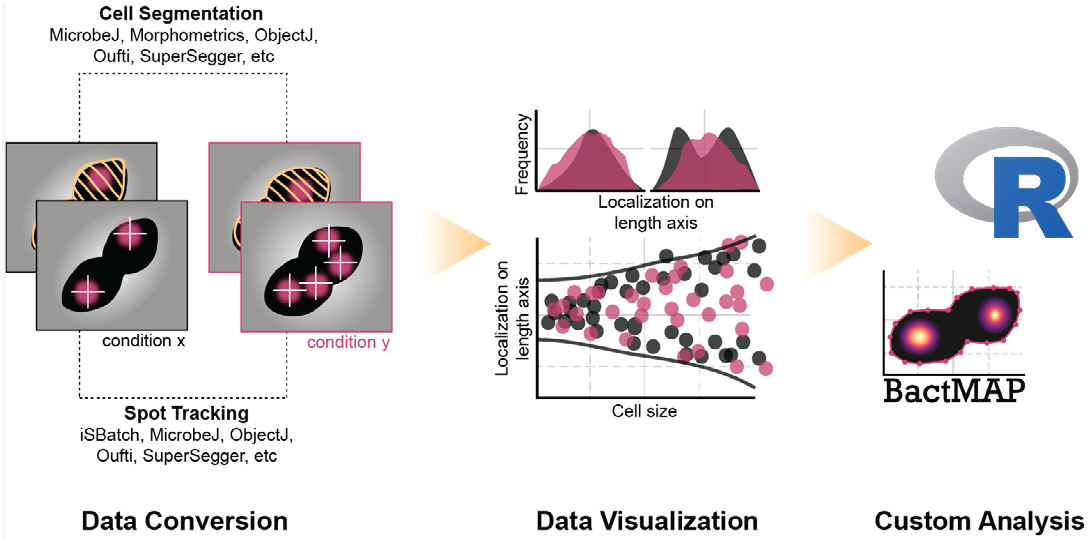
Visualization and analysis of microscopy data using BactMAP.

We benchmarked the various options of BactMAP by analyzing the localization of the replication fork and origin of replication in three different Gram-positive organisms: *Bacillus subtilis, Streptococcus pneumoniae* and *Staphylococcus aureus*. These organisms all have a different cell shape, leading to different challenges in segmentation. Indeed, we show that certain programs perform segmentation better on specific cell types than others (Fig S1 and S2). All data generated with these programs was transformed using BactMAP. To obtain useful biological insights from the data, the BactMAP toolbox was used to perform exploratory localization plotting, time-lapse analyses and finally we show how BactMAP can be used as a starting point for custom analysis with R.

## Results

### Visualization and analysis of microscopy data using BactMAP

BactMAP was built for automated import of segmentation data and for visualization of cells and their internal fluorescence signals (Fig. 1). To make BactMAP as useful as possible for the bacterial cell biology community, we standardized the import of datasets from five popular segmentation programs and one single-molecule tracking program: SuperSegger (Stylianidou *et al*., 2016), Microbej (Ducret *et al*., 2016), Oufti (Paintdakhi *et al*., 2016), Morphometrics (Ursell *et al*., 2017), ObjectJjChainTracer (Vischer *et al*., 2015; Syvertsson *et al*., 2016) and iSBatch (Caldas *et al*., 2015). Furthermore, new programs with improved or specialized cell segmentation capabilities are being developed constantly, as recently for instance BacStalk (Hartmann *et al*., 2018), a specialized tool for segmentation of cells with complex morphologies and the deep learning-based segmentation programs DeepCell (Bannon *et al*., 2018) and DeLTA (Lugagne et al. 2019). Therefore, to make BactMAP compatible with a wide range of current and future segmentation software packages, we also implemented a generic import function for segmentation input.

The visualization options of BactMAP are summarized in Fig. 2. In short, there are three main categories of plotting functions in BactMAP: 1) the visualization of intracellular fluorescence using information from the image itself, 2) visualization of subcellular localization of spots and objects as identified by any of the software packages and 3) plots which can be used in the analysis of time-lapse movies.

**Figure 2.**
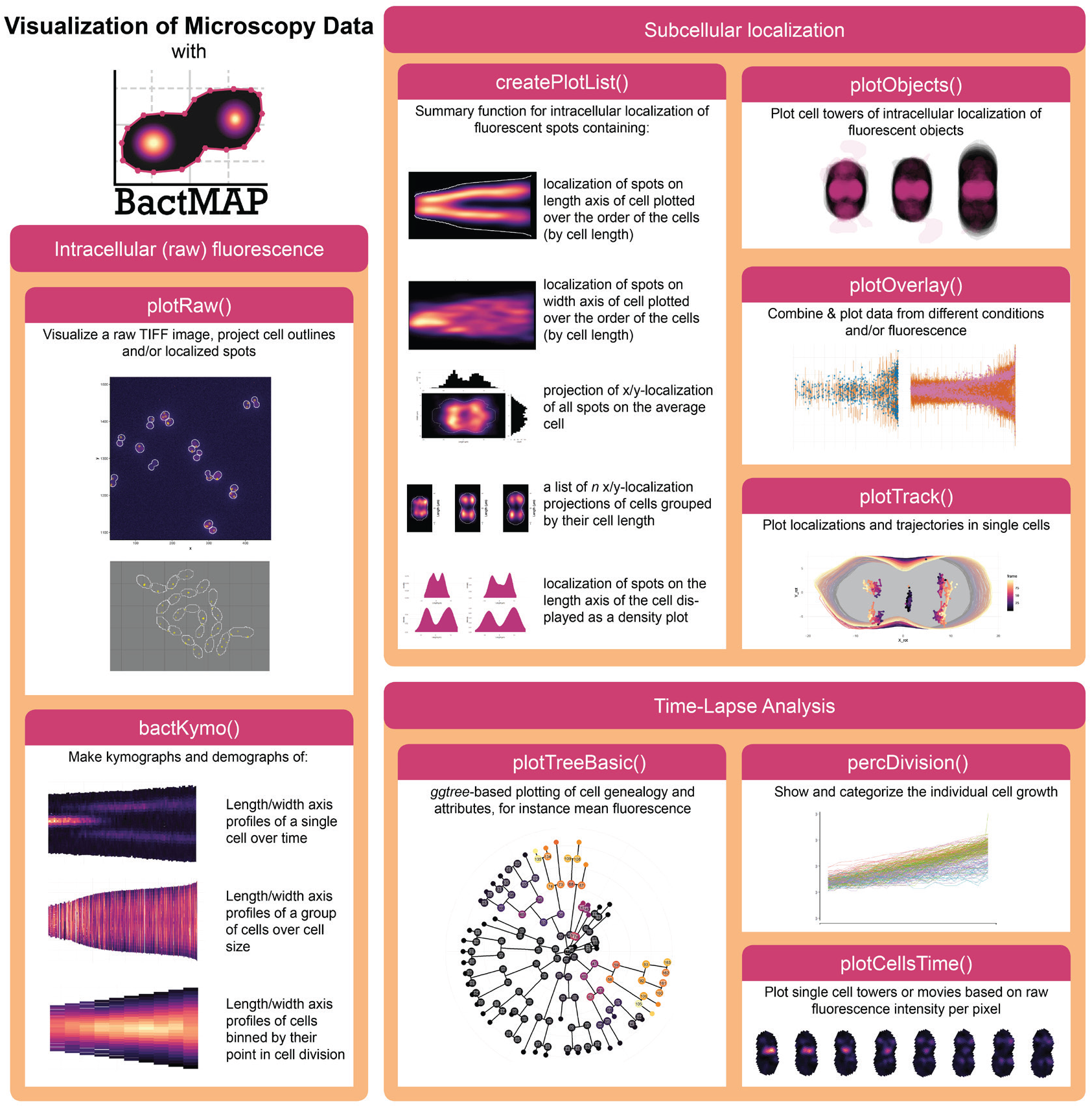
Overview of BactMAP’s plotting functions. Intracellular (raw) fluorescence. *plotRawQ* and *bactKymoQ* are both useful visualization tools which use cell outlines and the original image in TIFF format. *plotRawQ* shows the original microscopy pictures with the cellular outlines and/or the localization data. *bactKymoQ* makes kymographs and demographs of single cells and cell groups. **Subcellular localization.** For plotting of subcellular fluorescent spot localizations *createPlotlistQ* is used. This function returns a list of demographs, histograms an projections. For larger fluorescent objects, *plotObjectsQ* plots intracellular object shapes and localization through cell projections. When MicrobeJ or ISBatch are used to track fluorescent spots over time, *plotTrack()* can be used to visualize them. Moreover, *plotOverlayQ* can be used to plot cell towers and localization over time of different fluorescent channels and/or experimental conditions. Time-lapse analysis. *percDivisionQ* will categorize each cell based on growth speed and determine when a cell underwent a full division. *plotTreeBasicQ* uses the package *ggtree* (Yu et al. 2017) to plot Oufti’s or SuperSegger’s genealogy information as a tree plot. To visualize single-cell growth and fluorescence, *plotCellsTimeQ* uses cell outlines and raw microscopy images to create single-cell towers or movies.

### Exploratory localization plotting using BactMAP of the origin of replication in three differently shaped bacteria

To demonstrate the functionality of exploratory localization plotting using BactMAP, we analyzed the subcellular localization of the origin of replication in three Gram-positive bacteria: *Bacillus subtilis, Streptococcus pneumoniae* and *Staphylococcus aureus*. In *B. subtilis*, the origin of replication was visualized using a TetRjtetO system (*tetR-RFp, ori::tetD*, (Veening et al., 2009)), while in *S. pneumoniae* the origin was marked with a ParBp/*par*Sp-marker (*ParBp-GFp, ori.par-Sp*, (van Raaphorst *et al*., 2017)). In *S. aureus*, a protein-fusion of the native, origin-binding ParB to GFP was used as a marker for the origin of replication (Pinho and Errington, 2004). These three bacteria all have their own challenges in segmentation analysis due to differences in cell shape and mechanisms of growth and division.

*B. subtilis* divides by forming a cross-wall septum, which cannot be visualized by phase-contrast imaging. In this case, segmentation based on fluorescence images (e.g. cells stained with a fluorescent cell wall or membrane dye) is therefore necessary. The only currently available program which combines phase-contrast segmentation with semi-automated septum detection based on fluorescence is ChainTracer (Syvertsson *et al*., 2016). Other programs segment the *B. subtilis* chains well, but cannot locate septa. Morphometrics (Ursell *et al*., 2017) has an option for segmentation of peripheral fluorescence, which can be used for *B. subtilis* segmentation.

Encapsulated, oval-shaped *S. pneumoniae* cells can be difficult to segment, because they are small and form chains of cells under certain conditions (Domenech *et al*., 2018). As part of a BactMAP test session, we asked three of our lab members to use their program of choice to segment the same image of *S. pneumoniae* cells. The amount of cells detected by each program was comparable, however, measured cell widths and lengths were strikingly different between programs as well as between parameter settings (see Fig. S1). This underlines the importance of keeping the same globalsegmentation parameters within experiments, as well as of visual inspection of the segmentation results to make sure these are consistent. Since all programs are in principle capable of segmentation of phase contrast images of *S. pneumoniae* cells, the program choice largely depends on the goal of the experiment (e.g. comparing cell sizes, measuring internal fluorescence or tracking cells). To aid in this decision, we summarized the options and strengths of the five segmentation programs we tested in Fig. 3.

**Figure 3.**
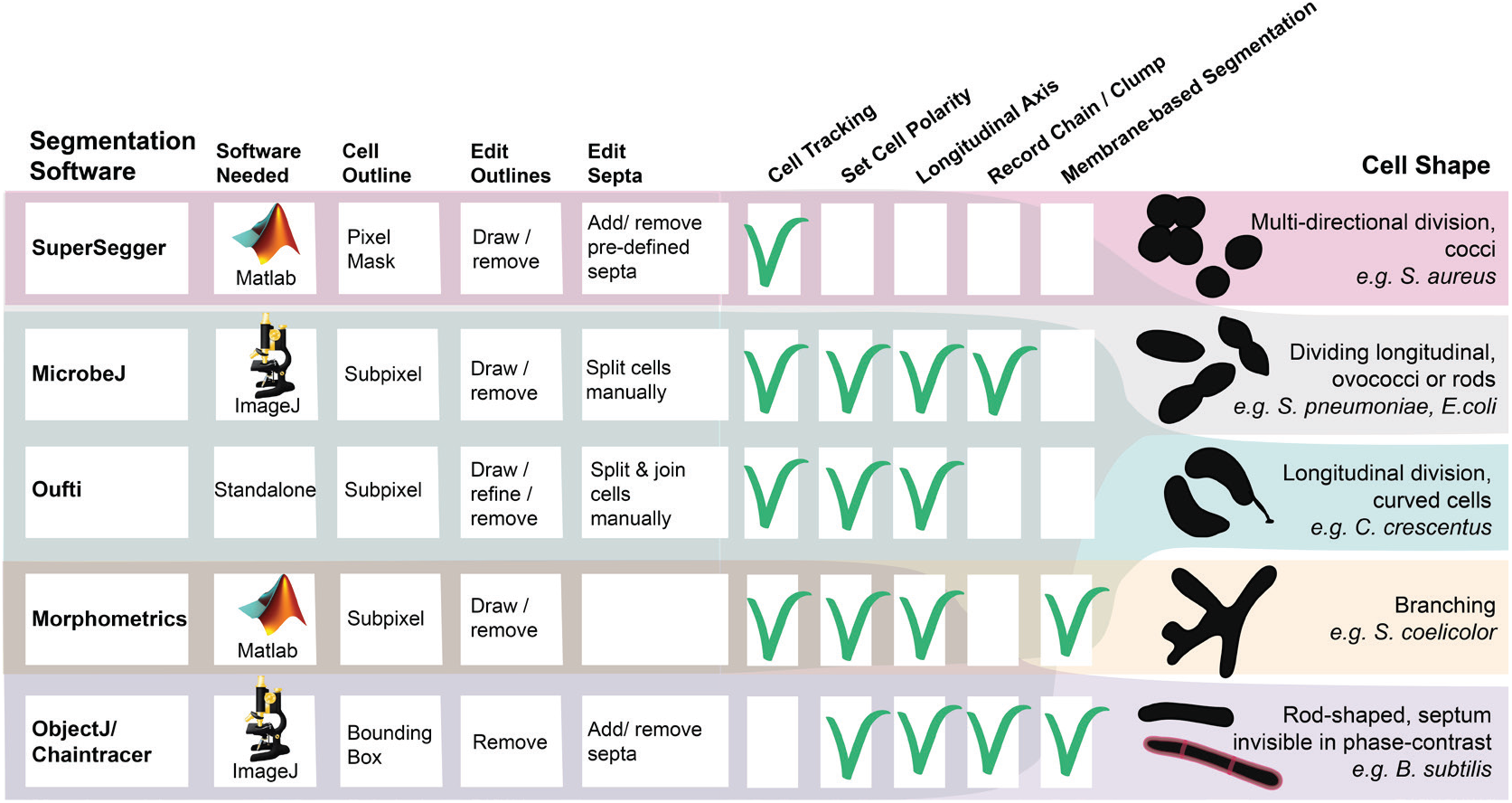
Overview of functionality of the five programs compatible with BactMAP. Of these, three are mat-lab-based (SuperSegger, Morphometrics and Oufti) and two are ImageJ Plugins (MicrobeJ, ObjectJ). While Oufti is Matlab-based, it comes as a standalone program for 64x operating systems. In addition to measuring the outlines, Oufti, MicrobeJ, Morphometrics and SuperSegger can track cells over time and provide information on growth speed and cell genealogy. Oufti, MicrobeJ, Morphometrics and ObjectJ estimate the cell length and curvature over the longitudinal axis. MicrobeJ offers a range of options for detection and counting of cell chains and clumps, while both MicrobeJ and ObjectJ offer options to detect cell features such as curvatures or invaginations as specified by the user. Finally, both SuperSegger and MicrobeJ give users the option to group cells based on user-specified cell features. All programs offer some options for manual editing of the results. In Oufti, a user can split or join cells, delete cells and draw new cell outlines. In Morphometrics, MicrobeJ and ObjectJ it is also possible to delete or add cells. For both Morphometrics and Oufti, it is not possible to move septa to a manually chosen subcellular location. In MicrobeJ this is possible, just as ObjectJ’s ChainTracer allows users to check, add and delete detected septa manually. Also in SuperSegger, it is possible to delete cells, but it is only possible to delete or add septa on pre-calculated positions.

*S. aureus* detection and segmentation is notoriously difficult because the cells are small cocci, not completely homogeneous in phase-contrast and they divide in multiple perpendicular planes (Pinho *et al*., 2013). For the analysis of *S. aureus* it was thus challenging to find a segmentation algorithm which created an outline of the whole cell based on phase contrast images. Both Oufti and ObjectJ are capable of creating a mask around *S. aureus* cells taken with phase-contrast microscopy, but both software packages have trouble attributing cell parameters to the mask because the algorithms are assuming a rod-shaped cell with a medial axis. Morphometrics and MicrobeJ both have the option of medial-axis based segmentation, and they also offer other options for subpixel segmentation, while SuperSegger only creates a pixel-resolution cell mask. We tested the last three programs and used BactMAP to display and compare the optimal segmentation result of the three programs (Fig. S2). All three programs performed well at segmenting the cells. Both MicrobeJ and Morphometrics, using a variety of different settings, however, still missed detection of many cells, while SuperSegger detected almost twice as many (Fig. S2). SuperSegger is therefore our program of choice for segmentation of *S. aureus* cells based on phase contrast images (see section below). Another option is to use fluorescence-based segmentation using a membrane dye or protein-fusion. Because Morphometrics has segmentation options for cells that are not rod-shaped, this would be the preferred option for fluorescence-based segmentation.

While different software packages were used for detection of cell outlines in the different species (as explained above), the single-molecule fluorescence detection program iSBatch (Caldas *et al*., 2015) was used to detect fluorescent foci for all three bacterial species (Fig. 4A). After imaging, fluorescence spot detection and cell detection, all datasets were imported using BactMAP.

**Figure 4.**
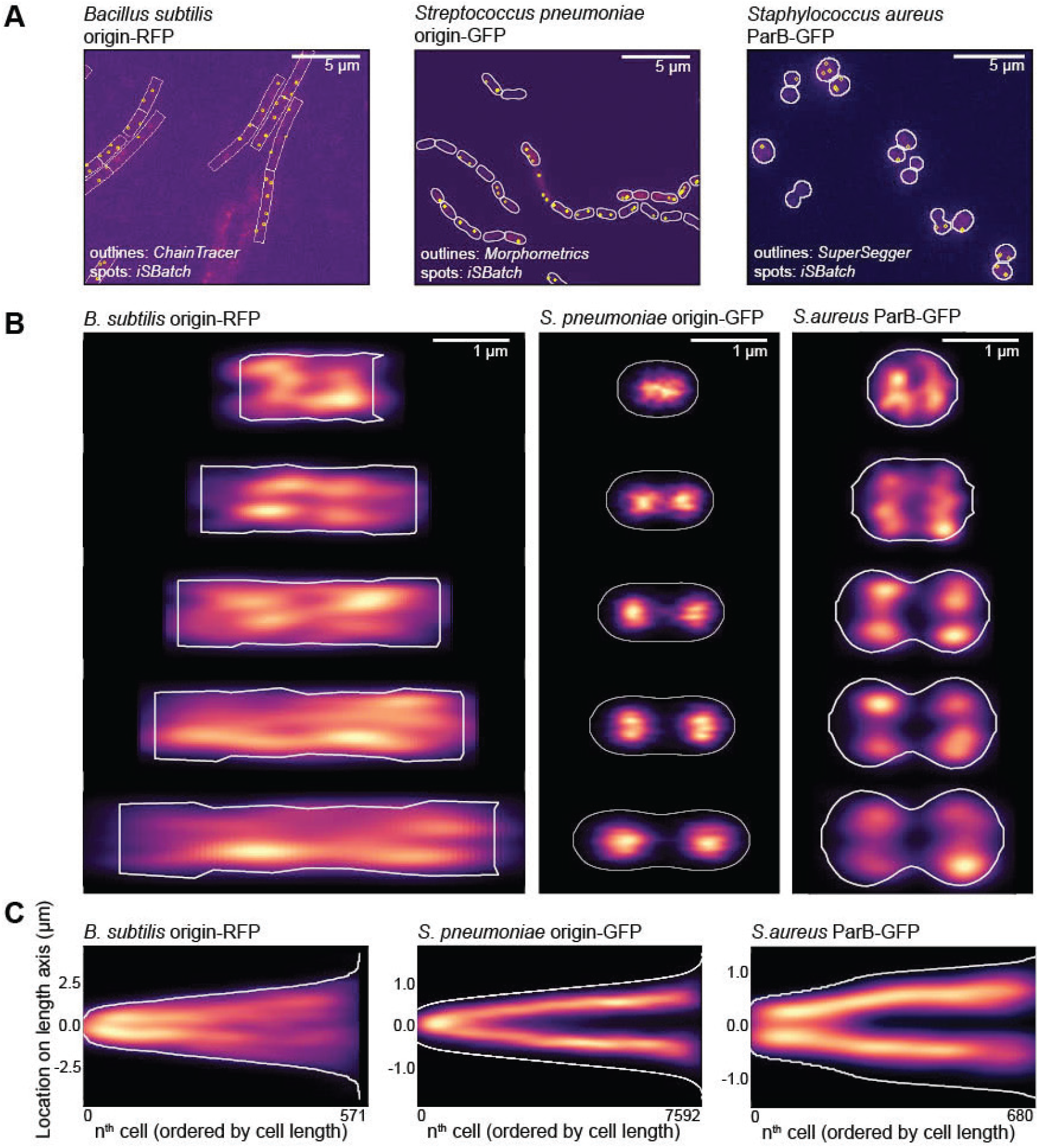
Exploratory plotting of the segmentation and origin localization of three differently shaped bacteria. **A.** For all three bacteria; a cutout of the raw image file with an overlay of the cell segmentation and detected fluorescent spots. The raw images, segmentation data and spot detection data was loaded into R using BactMAP’s extr.-functions, and the overlay images where created using *plotRawQ*. **B.** Cell towers showing the x,y-projection of the origin/parB inside the cell. The five groups are divided by cell length and contain an equal amount of cells. **C**. Projection of the localization of the origin/parB on the longest axis of the cell, where all cells are ordered by cell length.

Using BactMAPs *createPlotList*-function we plotted a projection of the subcellular localization of the origin of replication in five cell-size groups (Fig. 4B). *B. subtilis* was grown in lysogeny broth at 30°C. Due to multifork replication under these conditions, a median of 4 ± 2 origins/cell were detected (± indicating the standard deviation). The variable amount of spots lead to a diffuse signal in the density plots of the different cell size groups (Fig. 4B), but when only the coordinates of the origins on the length axis are plotted (Fig. 4C), a splitting of the origins between the two cell halves is visible. For *S. pneumoniae*, the origin of replication segregates early in the cell cycle, after which it moves to the future septa of the cell. In the dataset presented here, we detected 2 ± 2 spots/cell.

Similar to *S. pneumoniae, S. aureus* only initiates DNA replication one time per cell division. However, contrary to both *B. subtilis* and *S. pneumoniae, S. aureus* divides in consecutive perpendicular planes, making it especially hard to extract cell cycle information from microscopy snap shots. *S. aureus* cells only elongate slightly when the septal cross wall is being formed (Monteiro *et al*., 2015), after which they quickly split into two new daughter cells. To facilitate cell size grouping, we chose to record connected daughter cells as one cell (Fig. 4A and B, right panel). In this way, we could not only observe cells dividing in one plane, but also connected daughter cells that started to grow and divide in the perpendicular direction. Thus, in Fig. 4B, the two first cell groups represent cells before splitting, while the three last cell groups are cells already undergoing splitting. As observed in Fig. 4B, the average grouped cells indeed get mainly larger by length first, then by cell width. The origin is segregated in small cells, prior to splitting, without preference for location on the width axis (Fig. 4B). However, as observed in the third cell group, the second segregation takes place over the width axis (perpendicular to the initial segregation) and four spots are visible in the last three cell groups (Fig. 4B). This corresponds to the average amount of detected ParB-GFP spots in the *S. aureus* dataset (4 ± 3 spots/cell). In total, this analysis shows that BactMAP is capable of handling different input data and converting it into intuitive and qualitative as well as quantitative insightful reproductions of key cell cycle parameters.

### Single-cell time-lapse analysis of the replication fork of *S. pneumoniae* using BactMAP

In *S. pneumoniae*, the origin of replication moves to the new division sites very early in the cell cycle (Fig. 4B and C). The movement of the origin of replication is restricted, while the replication fork of *S. pneumoniae* is very mobile (van Raaphorst *et al*., 2017), in contrast to the replication fork of *E. coli* (Wallden *et al*., 2016) or *B. subtilis* (Liao *et al*., 2015, Mangiameli *et al*., 2018). Snap shots of fast-moving DnaX (the clamp loader of the replication fork) fused to GFP lead to blurred demographs, making it hard to interpret the localization of the replisome. Previously, we saw in single-cell movies using epifluorescence microscopy that the bulk of FtsZ and DnaX often localize on the same plane on the length axis of the cell. A recent study employing total internal reflection fluorescence (TIRF) microscopy (Perez *et al*., 2019) showed that FtsZ is not moving *en masse* from the division plane to the %-positions of the cell, but few treadmilling FtsZ clusters move along with MapZ, after which the concentration of FtsZ polymers colocalizing with MapZ gradually increases into the ring as seen by epifluorescence microscopy.

For our previous work (van Raaphorst *et al*., 2017), we imaged cells with DnaX-GFP and FtsZ-RFP (strain MK396) every 20 seconds for one hour. For this study, we re-analyzed these movies. We first used Oufti to track the cells and iSBatch to track internal FtsZ-RFP filaments and DnaX-GFP foci. After tracking of cells, filaments and foci, we used BactMAP to combine the information. Figure 5 displays some examples on how such data can be plotted with BactMAP. Figure 5A shows kymographs of the fluorescence intensity across the length-axis in a single cell over time. While the bulk of FtsZ has arrived at the %-positions of the cell around 10 minutes from the start of the movie, there is already an increase in fluorescence visible next to mid-cell from 4 minutes on. This is confirmed by the trajectories recorded by iSBatch (Figure 5B). Interestingly, the timing of movement from mid-cell to the quarter positions of the cell is not completely the same in the left and the right side of the cell, as was also shown by (Perez *et al*., 2019). As shown in Fig. 5C, the FtsZ-RFP tracks are short and sometimes even static. It is not surprising that we do not observe treadmilling in these conditions, since we imaged the cells in HiLo-mode (highly inclined and laminated optical sheet mode) where we focused on the middle of the cell and not in TIRF mode as performed by (Perez *et al*., 2019). While these conditions are good to follow DnaX and to see the general localization of FtsZ, we miss the focus on the edge of the cell and thereby the movement of FtsZ along the Z-ring.

**Figure 5.**
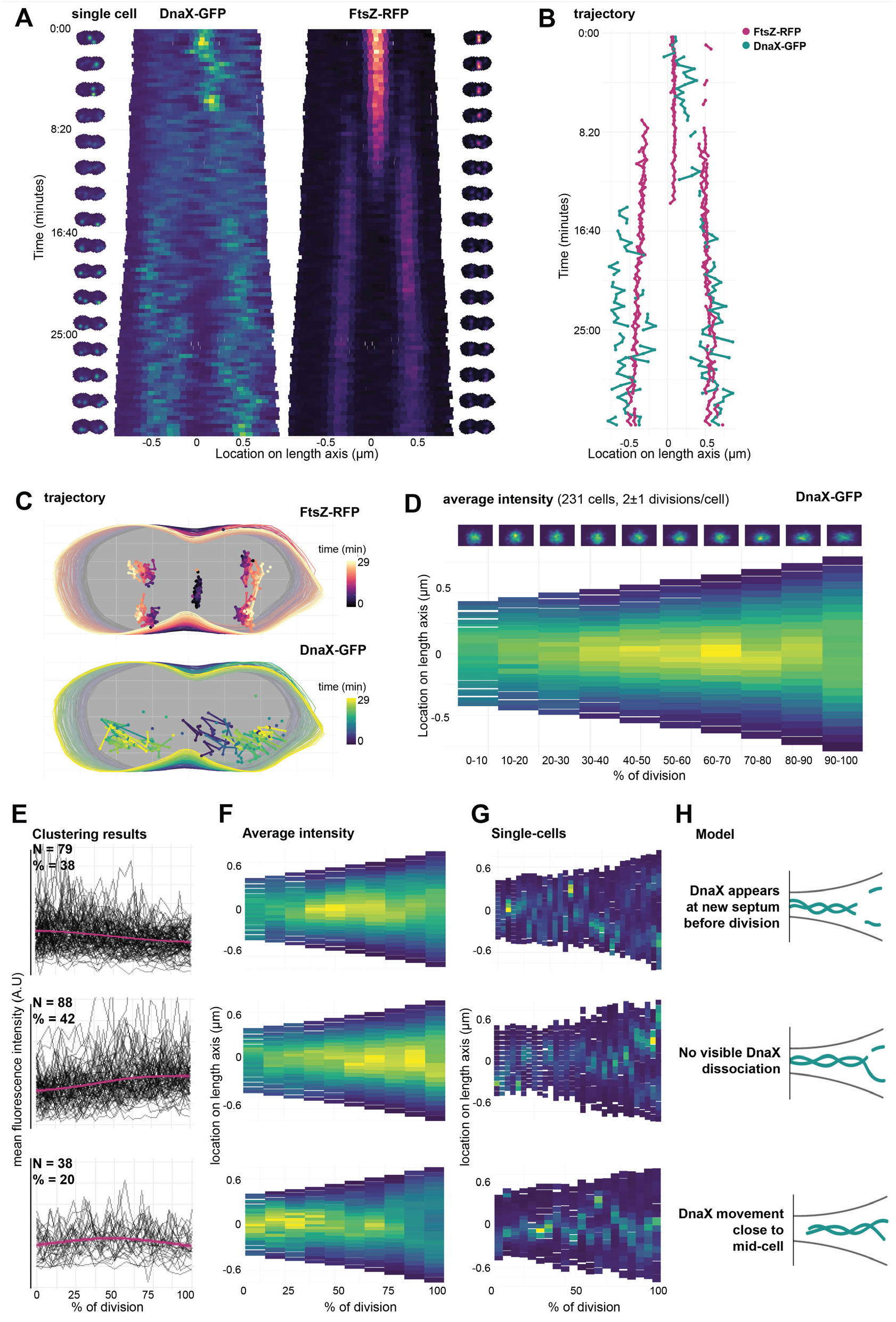
Single-cell timelapse analysis of the replication fork and FtsZ in *Streptococcus pneumoniae*. **A.** Kymographs of a single cell (strain MK396, *dnaX::dnaX-GFP-eryR, ftsZ::ftsZ-RFP-kanR)* imaged every 20 seconds for one hour. Cell outlines recorded with Oufti and combined with the raw image data using BactMAP. Left/right of the kymographs are movie strips of the single cell, created with BactMAP. **B.** Trajectory over the length-axis of the cell over time of FtsZ-RFP bundles and DnaX-GFP foci in the same cell as shown in A. Foci/bundles were tracked with iSBatch. **C.** X/Y trajectory over time of the cell shown in A and B. Outlines recorded with Oufti, tracks recorded with iSBatch. **D.** The growth curves of all cells were determined and curves of non-growing cells and cells with incomplete cell cycles were discarded. Cell parameters were binned in ten groups by % of division. Bottom: average intensity of GFP signal over the length axis of the cell per % of division. Top: density plots of X/Y coordinates of recorded GFP foci per binned % of division. X/Y coordinates were recorded with iSBatch, cell outlines with Oufti and the raw image files were used by BactMAP to determine the average intensity per bin. **E.** Clustering results: average cellular intensity over division percentage. Mean cellular fluorescence intensity (arbitrary units) over percentage of division (pink, standard deviation in shade), with single-cell fluorescence intensity paths shown in grey. Top-bottom shows each cluster, the amount of clusters (N) and the percentage of the total amount of cells (%). **F.** Average intensity profile. Average intensity of the length axis of the cell binned in 10 groups based on percentage of division for each of the three clusters (top-bottom). **G.** Single cells. Example kymographs of single cell members of each cluster. **H.** Schematic models of the DnaX dynamics.

While FtsZ moves to the new septa early during the cell cycle, DnaX seems to disassociate at mid-cell and assemble at the new quarter positions within a timeframe of 8 minutes. To see if we can observe association and dissociation in more cells, we re-analyzed a time-lapse movie of pneumococcus with DnaX-GFP imaged every 2 minutes for 4 hours (van Raaphorst *et al*., 2017). The lower time resolution allowed us to record multiple divisions while keeping the time resolution high enough to be able to see DnaX appear and disappear in single cells. BactMAPs function *perc_Division()* uses cell length as a proxy to determine when a cell in a time-lapse has divided, if this is not yet determined by the segmentation software used as input. After this, it uses the division time to determine how far into the cell cycle every single cell is at a given point in time. In this way, BactMAP can order and compare cells. Figure 5D shows a demograph of the fluorescence intensity over the length axis of the cell averaged by the percentage of division. Notably, no drop in fluorescence intensity is visible during cell division in this demograph, meaning that at the population level, DnaX association and dissociation is not measurable. This can be due to the lower time-resolution, difference in fluorescence intensity per cell or a high variability in association/dissociation timing between cells. This exploratory analysis shows that to follow the timing and localization of DnaX accurately, one needs to examine individual cells in time.

### Timing and localization of DnaX association/dissociation at the replication fork: custom analysis

Visual inspection of kymographs of individual cells over time showed that there is indeed a high variability in the localization of DnaX between cells. Using the R package TSclust(Montero and Vilar, 2014), we created a dissimilarity matrix and performed hierarchical clustering based on the fluorescence profile over the course of division of each cell (see Fig. S3). Before clustering, 10% of the cells were discarded because they showed no growth or too little fluorescence. After the clustering analysis, we grouped the cells into three main groups (Fig. 5E-H). In 38% of the remaining cells, the mean fluorescence drops just before division, after which it slightly increases again. This corresponds to cells where DnaX moves as in Fig. 5A-C: after moving around mid-cell, DnaX disappears for a few minutes, after which it reappears at the two new septa. However, in 42% of the cells, we observed no drop in fluorescence: DnaX moves gradually to the two cell poles. It could be that for this cell group, dissociation and association happened within our 2 minute interval. In 20% of the cells, DnaX remains at the old septum until division (Fig. 5E-H). It might be possible that the segmentation software split these cells earlier in their division cycle than the cells of the largest group, leading to an early drop in fluorescence instead of a late one. Segmenting cells based on membrane or cell wall markers could give more insight in whether this is the case. Nevertheless, this analysis using BactMAP provides new insights into DnaX dynamics during the pneumococcal cell cycle. We now show that the replication forks dissociate and associate before the cell splits and after the Z-ring is formed at the new ¼ positions.

## Discussion

In this work, we have developed BactMAP, a tool for visualization of microscopy data obtained from different image analysis programs. Using BactMAP, cell segmentation, fluorescence intensity and fluorescence spot detection results obtained from different software packages are all converted to the same format and can be plotted the same way. This allows for easy comparison of results by eye or by quantitative measures. BactMAP thus enables users to not only compare outputs from different software packages, but also combine them in one analysis. Finally, data from different experiments and different color channels can be easily combined in a uniform manner.

When creating BactMAP, our focus was to make a tool for plotting of intracellular fluorescence and automated import of different datasets. However, there are more types of analyses possible, some of which we slightly touched; for instance single molecule tracking, cell shape classification, analysis of cell lineages or visualization of localization microscopy. For advanced single molecule tracking, there are already software packages available, which can combine tracking analysis with single cell outlines from other programs (e.g. ISBatch (Caldas *et al*., 2015) and SMTracker (Rösch *et al*., 2018).

One of the more alarming outcomes of our analysis of segmentation outputs is that both the software and the user influence the outcome of the analysis to a great extent. Especially when comparing cell sizes from different experiments or when switching software within a lab, it is important to keep these limitations in mind and document the differences.

Using our newly developed R package BactMAP, we could revisit our previously analyzed microscopy images and time-lapse movies and gather new information. Most notably, using averaged datasets and observations of single-cell movies of DnaX-GFP movement during cell division, we concluded previously that the formation of the new replisome happens simultaneously with Z-ring formation. Using single-cell analysis instead of looking at the cells in population level, we see that the situation is more nuanced. In 38% of the cases, the replisome is formed after Z-ring formation, but in the remaining 62% of the cells we see DnaX foci moving gradually to the new septum or not moving away from the septum at all. In many cells, association and dissociation of the replication forks probably happens within a very short time interval. We also could get a more detailed view of global FtsZ movement during cell division.

Recently, more R packages and Shiny apps have been developed to enable researchers with little experience in R to use the plotting tools available with little effort, for example PlotsOfData (Postma and Goedhart, 2019). With BactMAP, researchers can use advanced plotting of microscopy data in R and have autonomy over both the way of plotting as well as the layout presented. After plotting, the data output can be used further for custom analysis. All in all, BactMAP is a tool for both initial data investigation and publication-ready plotting, and as a starting point for more in-depth data analysis. A user guide and the source code of BactMAP is available at https://veeninglab.com/bactmap.

## Experimental procedures

### Conversion to standard data structure

BactMAP converts data from segmentation and localization analysis tools into R dataframes with a standard structure, depending on the information enclosed in the input data: 1) meshframe, containing cell contours, 2) spotfTame, containing spot x/y coordinates, 3) objectframe, with x/y coordinates of the outline of fluorescent objects, 4) spots_relative and object_relative, with the x/y coordinates of the spots or object translated to intercellular coordinates, 5) timelapse-data, containing information on cell genealogy, and 6) cell-List, where the original input data is saved with as little change as possible.

BactMAP’s import functions are at least compatible with the current versions of segmentation and fluorescence detection sofware: Oufti (2015-current), Morphometrics (>version 0.1, Oct 25 2016), ObjectJ (version 1.04), MicrobeJ (version >5.13), SuperSegger (version 1.0.1) and isBatch (version >0.3.5).

Segmentation data standardly creates a meshframe, containing x/y-coordinates of the contours of the cells, the length and width of the minimal bounding box, cell and frame ID and the coordinates of the cell when the cells are turned on their long axis with their mid-point at [0,0]. The R package shotGroups (https://CRAN.R-project.org/package=shot-Groups) was used to create the function of finding the minimal bounding box and the cell angle needed to turn every cell on their length axis. For importing MATLAB (.mat) files directly into R, the R package R.matlab (https://CRAN.R-project.org/package=R.matlab) was used.

Similarly, when localization information is transformed by BactMAP, a spotframe is created which contains the coordinates of the spots and the frame ID. When segmentation and localization data are imported together, the data is combined to finally create the dataframe spots_relative. The R package SMDTools (VanDerWal *et al*., 2014) was used to build the function to connect the spot data to the cell outline data.

While the spotframe, meshframe and spots_relative data-frames are the most common output formats of transformation of BactMAP data, BactMAP also has standard functions to transform fluorescent object coordinates, cell genealogy information and raw TIFF microscopy images. A summary of transformation functions and their output can be found in the documentation of the BactMAP package available at https://veeninglab.com/bactmap.

### Visualization

The BactMAP plotting toolbox can be divided in three groups (see Fig. 2): (1) visualization of cells based on the raw image data and unprocessed cell outlines, (2) plots used to visualize, combine and categorize information on subcellular localization and (3) plots intended for time-lapse analysis (e.g. genealogy trees and extraction of single-cell movies).

One of the primary goals of this project was to make it easy to generate and edit publication-ready plots. For this reason, all plots generated using the BactMAP package are made using ggplot2 (Wickham, 2016). Since ggplots are easy to edit and build upon by adding layers of layouts, it is possible to change the color schemes and fonts or add new layers of data. All plotting functions also return the data used to make the plot, so a user can also decide to build the plot from scratch. Plots generated in R can be saved in vector-format PDFs. The standard color schemes used in BactMAP are colorblind-friendly (the viridis diverging heatmap color schemes or the palette designed by (Ichihara *et al*., 2010)). Using the R package gg-animate (https://gganimate.com/) both time-lapse and localization plots faceted by cell size can easily be converted into movies. When indicated, BactMAP plot outputs are compatible with the interactive plotting package plotly (https://plot.ly), which enables a user to change the axes manually, zoom by dragging/dropping or make interactive movies.

### Tutorials & Documentation

The source code and full documentation of all functions included in BactMAP can be found on https://veeninglab.com/bactmap, on the GitHub page of the Veening Lab (https://github.com/veeninglab), and are included in the BactMAP package.

The datasets used in this study can be downloaded as example datasetsfrom https://veeninglab.com/bactmap. Consequently, in a set of tutorials we explain how to recreate these plots using the example data on https://veeninglab.com/bactmap.

### Growth Conditions

*Bacillus subtilis* was incubated in Lysogeny Broth (LB, Miller) and grown O/N shaking (220 rpm) at 37°C. The culture was diluted 200 times in fresh LB and allowed to grow for 2 hours at 30°C. The culture was induced with D-Xylose (1% w/v, Sig-ma-Aldrich) for 30 minutes before the cells were washed in PBS containing 0.5 ug/mL Mitotracker Far Red (Invitrogen). The cells were washed again in PBS and where immobilized on 1% agarose in PBS as described before (de Jong et al. 2011) for microscopy.

*S. aureus* SH1000 carrying pLOW-parB-m(sf)gfp was grown in BHI broth (Oxoid) containing 5 ng/ml erythromycin overnight. The culture was diluted 100 times in fresh medium containing 100 |iM IPTG and incubated for 2.5 hours. Cells were then immobilized on 1% agarose in PBS for microscopy.

*Streptococcus pneumoniae* with DnaX-GFP (strain VL369/ RR23) and DnaX-GFP/FtsZ-RFP (strain VL469/MK396] were grown as described before (van Raaphorst *et al*., 2017). *Streptococcus pneumoniae* strain VL451/MK359 was grown in C+Y medium recipe 2018 (Domenech *et al*., 2018) until an OD of 0.1 and diluted 100 times in fresh medium containing 0.1 |iM ZnCl2. The culture was grown up to OD 0.1 again and immobilized on 1% agarose in PBS as described before for microscopy (de Jong *et al*., 2011).

**Table 1.** Strains used in this study

| Strain | Genotype | Reference |
| --- | --- | --- |
| *Bacillus subtilis* |  |  |
| VL429/JW147 | sp168; *hutM::tetO*, *Km<sup>R</sup>*, *amyE::Pxyl-tetR-mCherry*, *Spec<sup>R</sup>*, *PdnaN-gfp-dnaN*, *Cm<sup>R</sup>* | (Veening *et al.*, 2009) |
| *Staphylococcus aureus* |  |  |
| IM67 | SH1000, pLOW-parB-m(sf)gfp | Laboratory collection |
| *Streptococcus pneumoniae* |  |  |
| VL369/RR23 | D39V; *dnaX::dnaX-m(sf)GFP*, *ery<sup>R</sup>* | (van Raaphorst *et al.*, 2017) |
| VL451/MK359 | D39V, $\Delta bgaA::P_{Zn}$-*tetR-mKate2*, *parB<sup>mut</sup>-gfp*; *tet<sup>R</sup>*, *comCDE*, *parS<sub>p</sub>*, *kan<sup>R</sup>*; *pulA*, *tetO48*, *spc<sup>R</sup>* | (van Raaphorst *et al.*, 2017) |
| VL469/MK396 | D39V, *ftsZ::ftsZ-mKate2*, *kan<sup>R</sup>*, *dnaX::dnaX-m(sf)GFP*, *ery<sup>R</sup>* | (van Raaphorst *et al.*, 2017) |

### Microscopy

The images of *B. subtilis* and *S. pneumoniae* VL451/MK359 were acquired on a Leica DMi8 microscope with a DFC9000 GT camera, a 100x/1.40 NA phase-contrast objective (Leica) and a Lumencor SpectraX light engine with the following filter settings: GFP: SpectraX-Quad cube (Chroma #89000) 470/24 excitation & 515/40 emission; RFP: SpectraX-Quad cube (Chroma #89000) 575/35 excitation & 595/40 emission; far-red (MitoTracker Red): Alexa 633 cube (Leica #11103136): 610/655 excitation & 620/720 emission.

S. aureus was imaged using a Zeiss AxioObserver with ZEN Blue software. Images were captured using an 0RCA-Flash4.0 V2 Digital CMOS camera (Hamamatsu Photonics), a 100x phase contrast objective. HPX 120 Illuminator (Zeiss) was used as a fluorescence light source. The following filter settings were used for GFP: filter set 38 HE (Zeiss) 470/40 excitation and 525/50 emission.

*Streptococcus pneumoniae* with DnaX-GFP (strain VL369/ RR23) and DnaX-GFP/FtsZ-RFP (strain VL469/MK396) were imaged as described before (van Raaphorst *et al*., 2017). In short, the cells were imaged on a Deltavision Elite microscope in HiLo mode using a 488 nm (GFP) and 568 (RFP] laser through a Quad cube (Chroma #89000), either every 20 seconds in 30°C for one hour or every two minutes in 37°C for three hours an 48 minutes.

### Image Analysis

Images where converted to TIFF files using FIJI (Schindelin *et al*., 2012). Cell segmentation and fluorescence detection/ segmentation was done with isBatch (Caldas *et al*., 2015), MicrobeJ (Ducret *et al*., 2016), SuperSegger (Ducret *et al*., 2016; Stylianidou *et al*., 2016), Morphometrics (Ursell et al. 2017), ObjectJ (Vischer *et al*., 2015), ChainTracer (Syvertsson *et al*., 2016) and Oufti (Paintdakhi *et al*., 2016) as indicated in the results section. The parameters used for each experiment are documented in the tutorials and documentation on https://veeninglab.com/bactmap.

For testing the segmentation of *Streptococcus pneumoniae*, we asked members of the Veening lab to segment one single microscopy image with their segmentation program of choice. Four lab members including one of the authors participated, of which one lab member performed segmentation twice, using two different programs, resulting in five datasets. These datasets where analyzed as described in the results section. A full transcript of the analysis including the datasets can be found as a tutorial on https://veeninglab.com/bactmap.

## Acknowledgements

We would like to thank Ine Myrbraten (Norwegian University of Life Sciences) for help with construction of the *S. aureus* strain. We would like to thank Clement Gallay, Lance Keller and Jun Kurushima for providing segmentation data to build the package and participating in the pneumococcus segmentation comparison test. We would like to thank Vincent de Bakker and Doran Pauka for reading through & correcting the R code. Work in the Veening lab is supported by the Swiss National Science Foundation (SNSF) (project grant 31003A_172861), a JPIAMR grant (40AR40_185533) from SNSF and ERC consolidator grant 771534-PneumoCaTChER. M. Kjos is supported by a the Research Council of Norway (RCN, project number 250976) and a JPIAMR grant from RCN (project number 296906).

