## Supplemental Figures 1-3 for "BactMAP: an R package for integrating, analyzing and visualizing bacterial microscopy data"

### Supporting Information

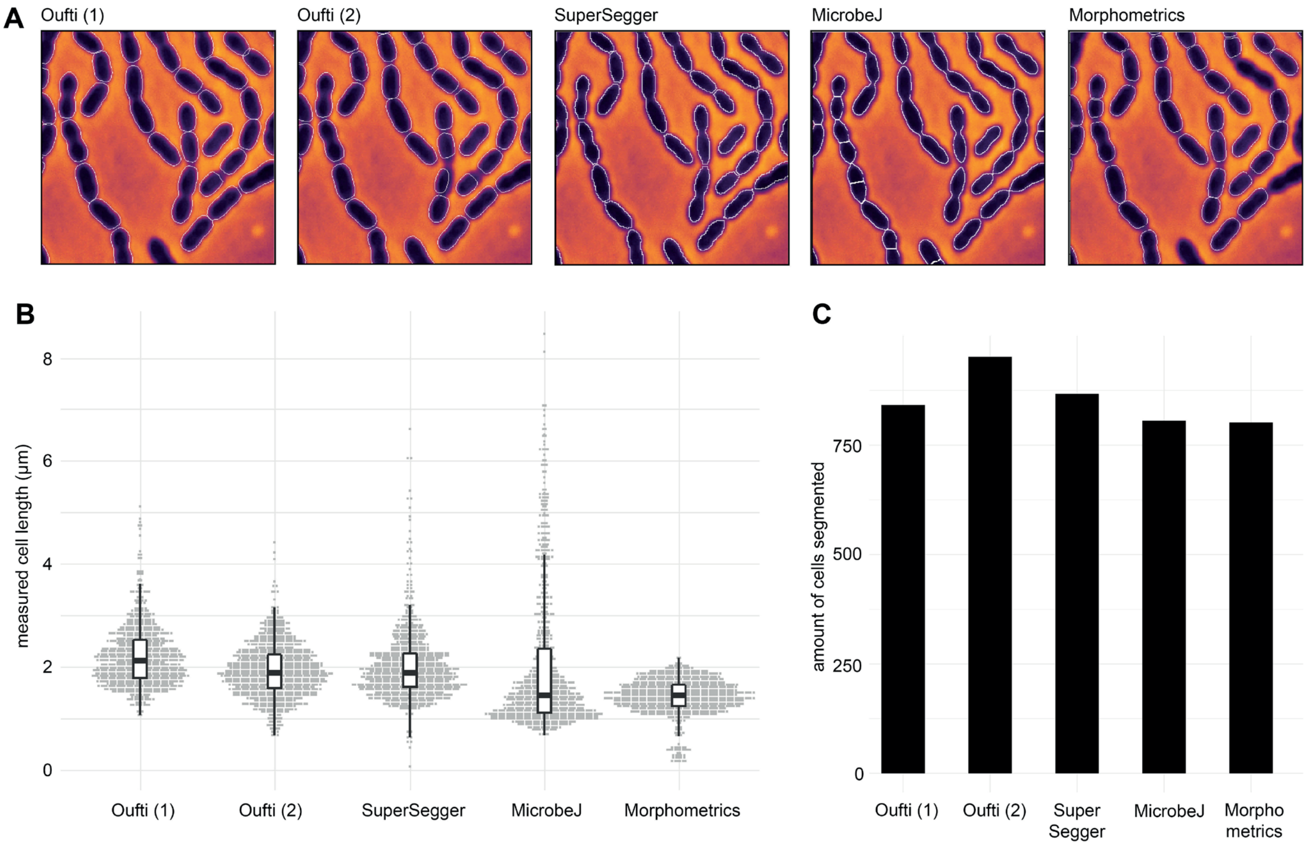


**Figure S1. Comparison of phase-contrast segmentation of *Streptococcus pneumoniae* cells by Oufti, Supersegger, MicrobeJ & Morphometrics.** Four lab members carried out cell segmentation of the same phase-contrast picture using the program they regularly use. One lab member decided to segment the image twice, one time using Oufti (Oufti (2) in the Fig. above) and one time using Supersegger. **A.** Cell outlines of a cutout of a microscopy image of chained *S. pneumoniae D39* cells (400*400 pixels or 20*20 µm) as segmented using the different programs. **B.** The measured cell sizes of all cells measured by each program. C: The amount of cells detected by each program.

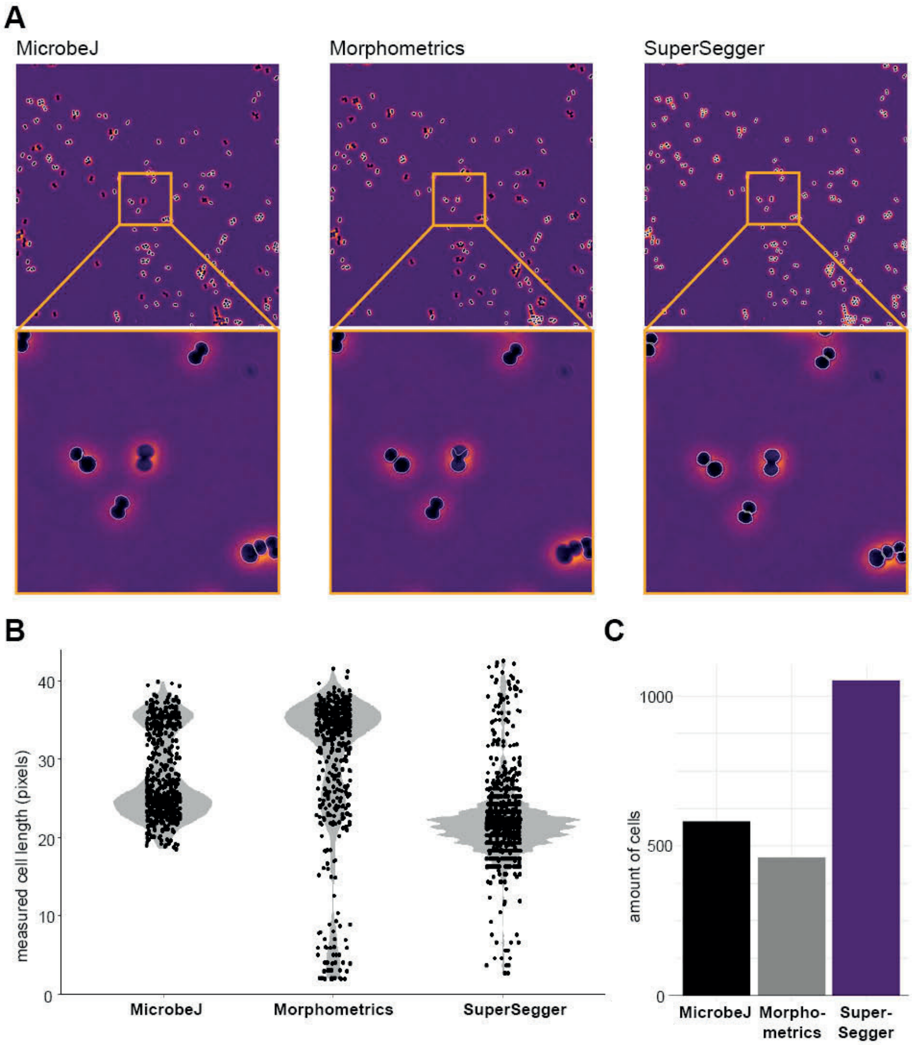


**Figure S2. Comparison of *Staphylococcus aureus* segmentation using MicrobeJ, Morphometrics and SuperSegger. A.** Plot of full phase-contrast image of *S. aureus* (2048*2048 $px^2$ or 113*113 $\mu m^2$) with cell outlines obtained with either MicrobeJ, Morphometrics or SuperSegger (left-right) with zoomed-in cutouts below (300*300 $px^2$ or 19.5*19.5 $\mu m^2$). **B.** Cell length distribution (in pixels) as measured by MicrobeJ, Morphometrics and SuperSegger, respectively. **C.** The total amount of cells found by MicrobeJ, Morphometrics and SuperSegger.

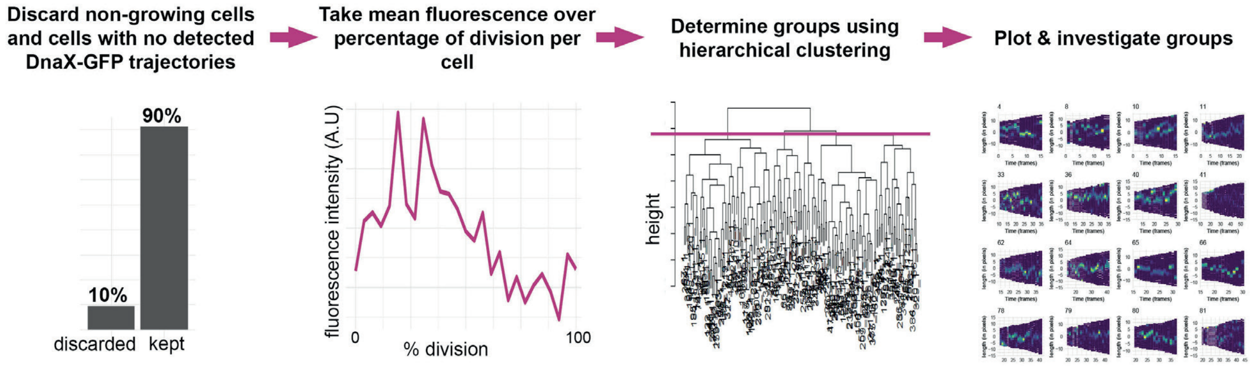


**Figure S3. Grouping cells by the behavior of DnaX-GFP over the course of division. Clustering workflow.** First, cells where discarded which did not show a full division, growth or fluorescence (26% of total). The mean fluorescence over time in each cell was determined, after which a dissimilarity matrix (based on correlation) was made and hierarchical clustering was performed. Using a cutoff of 3 groups, each group was investigated by plotting.
